# Structure of a transcribing RNA polymerase II–U1 snRNP complex

**DOI:** 10.1101/2020.10.19.344200

**Authors:** Suyang Zhang, Shintaro Aibara, Seychelle M. Vos, Dmitry E. Agafonov, Reinhard Lührmann, Patrick Cramer

## Abstract

To initiate co-transcriptional splicing, RNA polymerase II (Pol II) recruits U1 small nuclear ribonucleoprotein particle (U1 snRNP) to nascent pre-mRNA. Here we report the cryo-EM structure of a mammalian transcribing Pol II-U1 snRNP complex. The structure reveals that Pol II and U1 snRNP interact directly. This interaction positions the 5’ splice site in pre-mRNA near the RNA exit site of Pol II. Extension of pre-mRNA retains the 5’ splice site, leading to formation of an intron loop. Loop formation may facilitate scanning of the nascent pre-mRNA for the 3’ splice site and enable prespliceosome assembly and functional pairing of distant intron ends. Our results provide a starting point for a mechanistic analysis of co-transcriptional splicing and the biogenesis of mRNA isoforms during alternative splicing.

---

The production of mRNA in eukaryotic cells involves pre-mRNA synthesis and processing, in particular 5’ end capping, splicing, and 3’ end cleavage and polyadenylation. During splicing, non-coding introns are generally removed from the pre-mRNA in a co-transcriptional manner as the nascent RNA emerges from Pol II (*1–8*). Co-transcriptional splicing enhances the efficiency and accuracy of pre-mRNA processing and explains why splicing is at least ten times faster *in vivo* than *in vitro* (*9–11*). In metazoan cells, introns are often several thousand nucleotides long (*12*), posing the intriguing question how the ends of an intron are functionally paired for splicing. Co-transcriptional splicing was suggested to facilitate juxtaposition of the 5’ splice site (5’SS) and 3’SS (*13*). Consistent with this idea, the rate of Pol II elongation can affect selection of splice sites in pre-mRNA (*6–8, 14*), leading to alternative splicing and different mRNA isoforms (*15–17*). Pol II can recruit splicing factors via its flexible carboxyl-terminal domain (CTD) (*18–22*), which is however insufficient to stimulate splicing (*23*). Despite these advances, the mechanisms underlying co-transcriptional splicing remain unknown.

As a first step to investigate the mechanisms of co-transcriptional splicing, we studied the interaction between transcribing Pol II and U1 snRNP biochemically and structurally. U1 snRNP is the first spliceosomal building block to engage with nascent pre-mRNA and in human consists of U1 snRNA, seven Sm proteins and three U1-specific proteins, U1-70k, U1A and U1C (*24*). U1 snRNA recognizes the 5’SS through base-pairing (*25*). When the branch point sequence emerges on nascent pre-mRNA, U2 snRNP joins, forming the prespliceosome or A complex. The A complex later associates with the U4/U6.U5 tri-snRNP to form a fully assembled spliceosome, or pre-B complex (*26*), that is subsequently converted into the B complex and activated for splicing.

To assemble a transcribing Pol II-U1 snRNP complex, we used *S. scrofa* Pol II, which shows 99.9% sequence identity to human Pol II. We further used human U1 snRNP and a DNA-RNA scaffold that contains a DNA mismatch bubble and a modified MINX pre-mRNA with a 5’ cap (see materials and methods). The scaffold enables formation of a 9-base pair DNA-RNA hybrid duplex inside the bubble and contains a 145-nt (nucleotides) RNA that comprises a 5’ exon and a truncated intron of 29 nt (Fig. 1A and fig. S1, A and B). We incubated purified Pol II with the DNA-RNA scaffold and phosphorylated the resulting complex with the kinase positive transcription elongation factor b (P-TEFb). The phosphorylated complex was purified by size exclusion chromatography before its incubation with purified U1 snRNP (fig. S1C).

**Fig. 1.**
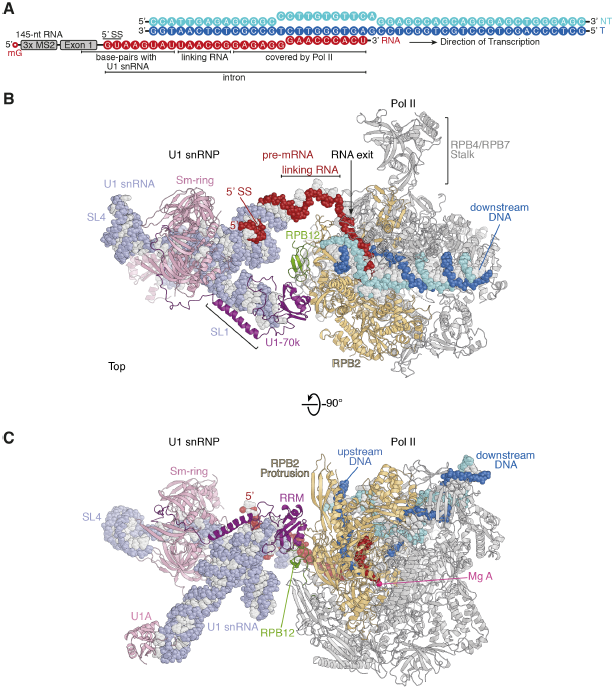
Structure of transcribing Pol II-U1 snRNP complex. (**A**) Nucleic acid scaffold with template DNA (T) in dark blue, non-template DNA (NT) in cyan and RNA in red. (**B** and **C**) Two views of the structure. Nucleic acids are shown in spheres. The backbone of U1 snRNA is in pale slate and U1 snRNP proteins are in pink, except for U1-70k in purple. Pol II subunits are in grey, except for RPB2 in gold and RPB12 in green. During transcription, Pol II moves to the right and RNA exits to the left. A magenta sphere depicts the Pol II active site. SL: stem loop, mG: 5’ cap.

The obtained sample was subjected to single-particle cryo-EM analysis (see materials and methods). A tilt angle of 40° was applied during data collection to resolve orientation bias (fig. S1, D to F, and table S1). Processing of the cryo-EM data showed that 17.5% of Pol II particles contained densities corresponding to U1 snRNP (fig. S2, A to F). Three-dimensional (3D) classification followed by 3D refinement of the class with the best density for U1 snRNP yielded a final reconstruction of the Pol II-U1 snRNP complex at an overall resolution of 3.6 Å, and focused 3D refinement improved densities for Pol II, the Pol II-U1 snRNP interface region, and U1 snRNP (fig. S2, G to I, figs. S3 and S4). Fitting and adjustment of the structures of Pol II (*27*) and U1 snRNP (*28, 29*) resulted in a model of the Pol II-U1 snRNP complex with very good stereochemistry (Fig. 1, B and C, table S1).

The structure shows that U1 snRNP binds directly to the surface of Pol II near upstream DNA and the site of RNA exit (Fig. 1). U1 snRNP engages Pol II via its conserved and functionally essential (*29*) subunit U1-70k, which contacts the protrusion domain in Pol II subunit RPB2 and the zinc finger domain in subunit RPB12 (Fig. 1, B and C, and Fig. 2A). U1-70k interacts with Pol II through its RNA recognition motif (RRM) domain that is resolved at ~3.5 Å resolution, revealing bulky side chains and enabling an unambiguous fit (fig. S3D and fig. S4, B and C). We did not observe any densities indicative of the Pol II CTD.

**Fig. 2.**
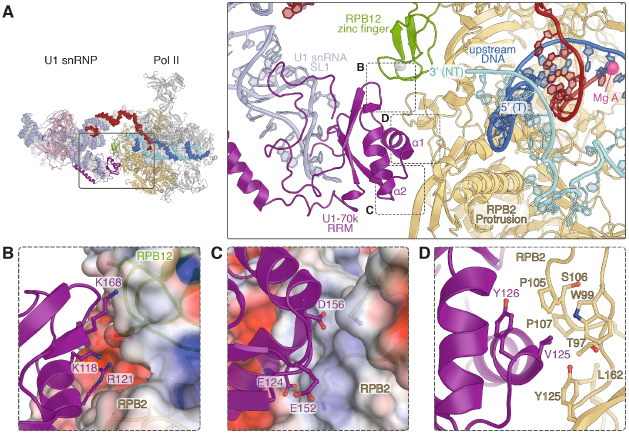
Direct Pol II-U1 snRNP interaction. (**A**) Close-up view of the Pol II-U1 snRNP interface showing interactions between the RRM domain of U1-70k (purple) and Pol II subunits RPB2 (gold) and RPB12 (green). The Pol II active site is depicted with a magenta sphere. (**B** to **D**) Detailed views of interactions between residues of U1-70k RRM and Pol II. The electrostatic surface potential displays a range of ± 5 kT/e, where blue and red represent positively and negatively charged areas, respectively.

Our structure provides details of the Pol II-U1 snRNP interface. The U1-70k RRM interacts with Pol II mainly through its two conserved a-helices (human residues 116-126; 154-164) (Fig. 2A). The Pol II-U1 snRNP interacting surfaces are reversely charged (fig. S5, A and B). The side chain of the positively charged U1-70k RRM residue Arg121 inserts into a negatively charged pocket formed by RPB2 and RPB12 (Fig. 2B). Three negatively charged residues (Glu124, Glu152 and Asp156) of U1-70k RRM contact a positively charged surface of the RPB2 protrusion domain (Fig. 2C). In addition, Val125 of U1-70k RRM binds to a hydrophobic patch of RPB2 (Fig. 2D).

Residues in U1-70k that contact Pol II are highly conserved among metazoa (fig. S5C). RPB2 residues that interact with U1-70k are also well conserved among multicellular eukaryotes and differ from those in the counterpart subunits of Pol I and Pol III (fig. S5D). This indicates that the observed U1 snRNP interaction with Pol II is conserved among metazoa and specific to the Pol II transcription system. In contrast, the interacting residues in both U1-70k and RPB2 are only partially conserved in yeasts (fig. S5, C and D), implying that the mechanism of co-transcriptional splicing may differ in some unicellular eukaryotes, which contain fewer and shorter introns.

Superposition of our Pol II-U1 snRNP structure with the activated Pol II elongation complex (*27*) EC* revealed that binding of U1 snRNP, as observed in our structure, is compatible with the presence of the transcription elongation factors DSIF, SPT6, and PAF1 complex (PAF) on the Pol II surface (fig. S6). Consistent with this compatibility, a stable complex containing EC* and U1 snRNP can be formed (Fig. 3, A to C, and fig. S7, A to F). In summary, these results reveal an unexpected direct Pol II-U1 snRNP interaction that is compatible with the presence of general elongation factors in the transcribing complex EC*.

**Fig. 3.**
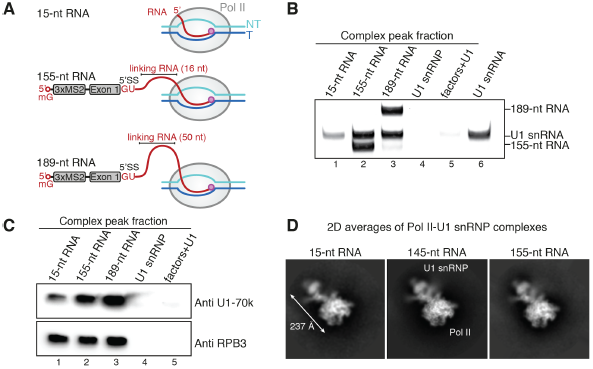
RNA-independent Pol II-U1 snRNP interaction. (**A**) Schematic of Pol II complexes with various RNAs. The active site of Pol II is depicted with a magenta sphere. (**B** and **C**) EC* associates with U1 snRNP in the presence of RNA lacking the 5’SS (15-nt) or RNAs with an extended linking region (155-nt and 189-nt). Shown is the analysis of the complex peak fraction (fraction #2 of fig. S7A) obtained by size exclusion chromatography using denaturing gel electrophoresis (B) or Western blotting (C). (**D**) Cryo-EM analysis of Pol II-U1 snRNP complexes containing 15-nt RNA or 155-nt RNA showed 2D averages resembling those of the refined structure with 145-nt RNA.

Our structure additionally shows one turn of an RNA duplex formed by base pairing of the 5’SS with U1 snRNA (Fig. 1 and fig. S4, A and D), as observed in previous U1 snRNP structures (*28–31*). The pre-mRNA thus tethers Pol II to U1 snRNP using a 6-nt long, flexible linking RNA region (Fig. 1A and fig. S4A). To investigate the role of RNA tethering, we performed analytical size exclusion assays using the activated elongation complex EC* (see materials and methods). We prepared EC* containing a minimal scaffold with a 15-nt RNA that lacks the 5’SS and is completely sequestered within the Pol II core (Fig. 3A and fig. S1, A and B), excluding any RNA-based contacts between Pol II and U1 snRNP. The resulting minimal EC* still bound U1 snRNP, although at apparently sub-stoichiometric levels (Fig. 3, B and C, and fig. S7, A and B).

To confirm that the direct Pol II-U1 snRNP interaction occurs in the absence of a tethering RNA, we subjected a Pol II-U1 snRNP complex containing the nucleic acid scaffold with the 15-nt RNA to cryo-EM analysis. We obtained the same two-dimensional particle averages as for our original structure (Fig. 3D). The percentage of Pol II-U1 snRNP particles was lower, hindering calculation of a 3D reconstruction. Nevertheless, these results show that the tethering RNA is not required for Pol II-U1 snRNP interaction *in vitro*, explaining why U1 snRNP can be recruited to transcription units independent of splicing *in vivo* (*32*).

Our observations suggested that transcription elongation may generate an RNA loop between Pol II and U1 snRNP while the direct Pol II-U1 snRNP interaction is maintained. To investigate this, we tested whether insertion of additional nucleotides into the linking RNA region still allows for U1 snRNP binding to Pol II or EC*. We designed scaffolds that contained either a 155-nt RNA that has a 10-nt extension in the linking RNA compared to the 145-nt RNA in the cryo-EM structure, or a 189-nt RNA that contains a linking RNA of 50 nt and would be sufficient for formation of an active spliceosomal B complex (*33*) (Fig. 3A and fig. S1, A and B). Both scaffolds with extended RNAs allowed formation of an EC*-U1 snRNP complex (Fig. 3, B and C, and fig. S7, A to D). These results support the model that the 5’SS is retained near the Pol II surface while the intron loops out as pre-mRNA is elongated.

To confirm that the same Pol II-U1 snRNP interaction is maintained when the linking RNA is extended, we determined the cryo-EM structure of the Pol II-U1 snRNP complex with the scaffold containing the 155-nt RNA at a nominal resolution of 3.9 Å (fig. S8). The structure was essentially unaltered compared to our first structure, revealing the same direct Pol II-U1 snRNP interaction (Fig. 3D and fig. S8). In addition, *in vitro* RNA elongation activities of Pol II or EC* were not affected by the presence of U1 snRNP (fig. S7, G and H). These results indicate that U1 snRNP can remain bound to Pol II while pre-mRNA is elongated during transcription.

To investigate whether assembly of spliceosomal complexes may be possible on the Pol II surface, we compared our structure with the yeast A complex (*34*) (fig. S9, A and B). This suggested that U2 snRNP is accommodated, indicating that the A complex can form on Pol II without moving U1 snRNP. Human A complex formation requires binding of stem loop 4 (SL4) of U1 snRNA to the U2 snRNP subunit SF3A1 (*35*). In our structure, SL4 faces away from Pol II and is available for SF3A1 interaction (Fig. 1, B and C). Superposition of our structure onto the human pre-B complex (*30, 31*) resulted in clashes between Pol II and the U4/U6.U5 tri-snRNP (fig. S9C). However, the tri-snRNP is flexibly attached to the remainder of the pre-B complex (*26*), and structural adjustments may allow formation of the pre-B complex on Pol II.

Our findings suggest a topological model for co-transcriptional splicing (Fig. 4). In this model, U1 snRNP is recruited when the 5’SS emerges in nascent pre-mRNA. A direct Pol II-U1 snRNP interaction is established, retaining the 5’SS near the RNA exit site of Pol II. This results in formation of an intron loop and facilitates scanning for the branch point and the 3’SS in nascent pre-mRNA. Recruitment of U2 snRNP then leads to formation of the A complex on the Pol II surface. Binding of the U4/U6.U5 tri-snRNP may allow formation of the pre-B complex on Pol II, but the subsequent transition to the B complex displaces U1 snRNP, thereby detaching the spliceosome from Pol II. This liberates Pol II and U1 snRNP for detection of the next 5’SS and repetition of the co-transcriptional splicing cycle on the next transcribed intron.

**Fig. 4.**
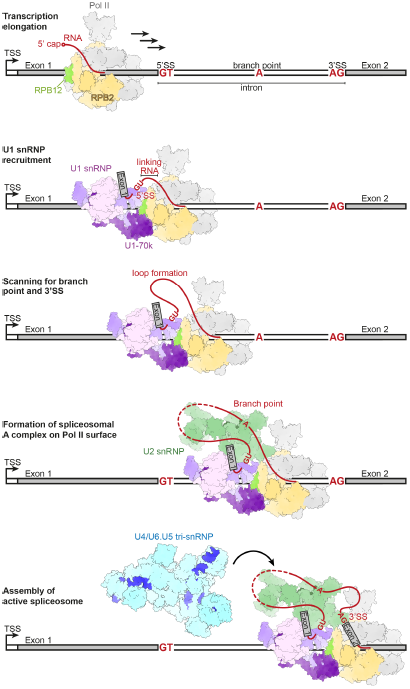
Model of co-transcriptional splicing. The intron is defined by the 5’SS, branch point and 3’SS (conserved nucleotides in red). When the 5’SS emerges in nascent pre-mRNA, U1 snRNP (purple) is recruited and directly binds Pol II subunits RPB2 (gold) and RPB12 (green). U1 snRNP and the 5’SS are retained near the RNA exit site during transcription elongation, resulting in the formation of an RNA loop. Loop formation facilitates scanning for the downstream branch point and 3’SS and assembly of the A complex.

Finally, U1 snRNP inhibits 3’ end processing (*36*), suppresses premature transcription termination (*37*), and impairs premature termination in the sense direction of bidirectional promoters (*38, 39*). We speculate that U1 snRNP binding to the Pol II surface interferes with productive binding of the 3’ end processing machinery. Thereby U1 snRNP may suppress premature termination and keep Pol II in an actively transcribing state until a genuine 3’ end processing signal is detected in nascent RNA.

## Acknowledgments

We thank C. Dienemann and U. Steuerwald for support at the microscope and maintaining the EM facility, T. Schulz for maintaining cells and thymus, and P. Rus and U. Neef for running the insect cell facility.

## Author contributions

S.Z. designed and performed all experiments, collected and analyzed cryo-EM data, determined the structure and carried out biochemical analysis. S.A. assisted with cryo-EM data processing. S.A. and S.M.V. provided advice. S.M.V. and D.E.A. performed related preliminary experiments. D.E.A. and R.L. provided U1 snRNP. R.L. provided advice. P.C. supervised research. S.Z. and P.C. wrote the manuscript, with input from R.L.

## Data and materials availability

The cryo-EM reconstructions and final model were deposited with the EMDB under accession code XXXX and the PDB under accession code XXXX. Correspondence and request of materials should be addressed to P.C.

## Materials and Methods

### Expression and protein purifications

The porcine Pol II was purified from *S. scrofa* thymus as described (*27*). In brief, the thymus was flash frozen in liquid nitrogen and homogenized in a blender. After removing the fat tissue, the supernatant was precipitated with 0.04% polyethylenimine (PEI). The PEI pellet was dissolved in 0.4 M ammonium sulfate buffer and applied onto a column packed with Macro-Prep High Q resin (Biorad). The eluted peak was then precipitated with 4 M ammonium sulfate and the re-dissolved pellet applied onto an 8WG16 antibody column. The eluate was further purified on a Uno Q1 column (Biorad) with a final size exclusion purification step using the Sephacryl S-300 16/60 column (GE Healthcare) in 10 mM HEPES pH 7.25, 150 mM NaCl, 10 μM ZnCl2 and 10 mM DTT. Human U1 snRNP was purified from HeLa cell nuclear extract by immunoaffinity purification and a 10-30% glycerol gradient as described (*28*). The fractions containing U1 snRNP were diluted to 100 mM NaCl and applied onto a heparin sepharose column (GE Healthcare). U1 snRNP was eluted with 20 mM Tris-HCl pH 7.9, 450 mM NaCl, 1.5 mM MgCl_2_ and 1 mM DTT. Human P-TEFb, PAF, DSIF and SPT6 were purified as described (*27*).

### RNA preparation

The following RNA constructs were generated for this study:

15-nt RNA: 5’-*GAG AGG GAA CCC ACU*-3’

145-nt RNA: 5’-3xMS_2_-CUU GGA UCG GAA ACC CGU CGG CCU CCG **ACA GGU AAG UAU** UAA CCG *GAG AGG GAA CCC ACU*-3’

155-nt RNA: 5’-3xMS_2_-CUU GGA UCG GAA ACC CGU CGG CCU CCG **ACA GGU AAG UAU** CUA GCA UGU AUA ACC G*GA GAG GGA ACC CAC U*-3’

189-nt RNA: 5’-3xMS_2_-CUU GGA UCG GAA ACC CGU CGG CCU CCG **ACA GGU AAG UAU** CUA GCA UGU AGA ACU GGU UAC CUG CAG CCC AAG CUU GCU GCA CGU AAC CG*G AGA GGG AAC CCA CU*-3’

3xMS_2_: 5’-CGU ACA CCA UCA GGG UAC GAG CUA GCC CAU GGC GUA CAC CAU CAG GGU ACG ACU AGU AGA UCU CGU ACA CCA UCA GGG UAC G-3’

The sequence that is complementary to U1 snRNA is in bold and the sequence that is covered within Pol II is in italic. The 15-nt RNA was synthesized by IDT. All other constructs contained modified MINX pre-mRNA sequence (*40*) and were cloned into the pUC18 vector with a T7 promotor and three MS_2_-tags at the 5’ end and an HDV (hepatitis delta virus) ribozyme at the 3’ end. The plasmids were prepared using the Maxiprep kit (Qiagen) and digested with XbaI overnight. The digested plasmids were purified by phenol chloroform extraction and ethanol precipitation. RNA was prepared by *in vitro* transcription using T7 RNA polymerase (5 U/μl final, Thermo Fisher Scientific) at 37 °C for 4 hours in the presence of 100 ng/μl digested plasmid, 40 mM Tris-HCl pH 7.9, 30 mM MgCl_2_, 10 mM DTT, 10 mM NaCl, 2 mM spermidine, 4 mM NTP and 0.001% Triton X-100. The transcription reaction was incubated at 60 °C for 20 min to facilitate HDV ribozyme cleavage, and then precipitated with ethanol, dissolved in 1x RNA loading dye (47.5% formamide, 0.01% bromophenol blue and 0.5 mM EDTA) and loaded onto 6% denaturing gels (6% acrylamide/bis-acrylamide, 7M urea and 1x TBE). The RNA bands were excised from the gels using UV shadowing and incubated in 0.3 M sodium acetate pH 5.2 overnight at −80 °C for extraction. The RNA was subsequently precipitated with isopropanol at −80 °C for 1 hour and dissolved in water.

RNAs were 5’ capped using the Vaccinia capping system (NEB). For each reaction of 20 μl, the RNA (10 μg) and nuclease-free water were combined to a final volume of 15 μl, heated at 65 °C for 5 min, put on ice for 5 min and the following components were added in the specified order: 1x capping buffer (NEB), 0.5 mM GTP, 0.1 mM SAM and 1 μl Vaccinia capping enzyme (10 units/μl, NEB). The reaction was incubated at 37 °C for 1 h and the capped RNA was purified by phenol chloroform extraction and ethanol precipitation at −20 °C overnight. The precipitated RNA was dissolved in water and flash frozen in small aliquots in liquid nitrogen.

### Sample preparation for cryo-EM

The transcribing Pol II-U1 snRNP complex was formed on a DNA scaffold with the following sequences: template DNA 5’-/Desthiobiotin/-GCT CCC AGC TCC CTG CTG GCT CCG AGT GGG TTC TGC CGC TCT CAA TGG-3’, non-template DNA 5’-CCA TTG AGA GCG GCC CTT GTG TTC AGG AGC CAG CAG GGA GCT GGG AGC-3’. The DNA scaffold contains 14 nucleotides of upstream DNA, 23 nucleotides of downstream DNA and a mismatch bubble of 11 nucleotides, of which 9 nucleotides of the template DNA base-pairs with the RNA. The DNA oligos were synthesized by IDT and dissolved in water.

RNA (15-nt, 145-nt or 155-nt) and template DNA were mixed in equimolar ratio (30 μM) in 20 mM HEPES pH 7.4, 100 mM NaCl and 3 mM MgCl_2_, and annealed by heating up at 60 °C for 4 min followed by decreasing the temperature by 1 °C min^-1^ steps to a final temperature of 30 °C in a thermocycler. *S. scrofa* Pol II (50 pmol) and the RNA–template hybrid (100 pmol) were incubated for 10 min at 30 °C, followed by addition of the non-template DNA (200 pmol) and incubation at 30 °C for another 10 min. Phosphorylation of Pol II was performed with 0.4 μM P-TEFb and 1 mM ATP in 20 mM HEPES pH 7.4, 100 mM NaCl, 3 mM MgCl_2_ and 1 mM DTT in a total volume of 25 μl for 30 min at 30 °C. The assembled transcribing Pol II complex was then applied onto a Superdex 200 Increase 3.2/300 column (GE Healthcare) equilibrated with the SEC150 buffer (20 mM HEPES pH 7.4, 150 mM NaCl, 3 mM MgCl_2_, 0.5 mM tris(2-carboxyethyl)phosphine (TCEP)). Purified U1 snRNP from the heparin column was further purified on the Superose 6 Increase 3.2/300 column (GE Healthcare) in SEC150 buffer. The peak fraction of the Pol II complex was mixed with the peak fraction of U1 snRNP at a molar ratio of 1:1.5, incubated on ice for 30 min and used for freezing grids. Similar results were obtained when combining the Pol II complex and U1 snRNP before the size exclusion purification step.

Samples were diluted to a concentration of 60-65 nM for the Pol II-U1 snRNP complex and 2 μl was applied to each side of the R2/2 UltrAuFoil grids (Quantifoil) that had been glow-discharged for 100 s. After incubation of 10 s and blotting for 4 s, the grid was vitrified by plunging it into liquid ethane with a Vitrobot Mark IV (FEI Company) operated at 4 °C and 100% humidity.

### Analytical size exclusion assay

Analytical size exclusion assay was performed using the same DNA scaffold as for the cryo-EM studies. EC* had to be used instead of Pol II to allow separation of the U1 snRNP bound complex from free U1 snRNP. The same procedure was carried out for three different RNA constructs (15-nt, 155-nt and 189-nt RNAs). RNA and template DNA were mixed in equimolar ratio (25 μM) in 20 mM HEPES pH 7.4, 100 mM NaCl and 3 mM MgCl_2_, and annealed by heating up at 60 °C for 4 min followed by decreasing the temperature by 1 °C min^-1^ steps to a final temperature of 30 °C in a thermocycler. *S. scrofa* Pol II (25 pmol) and the RNA–template hybrid (30 pmol) were incubated for 10 min at 30 °C, followed by addition of the non-template DNA (50 pmol) and incubation at 30 °C for another 10 min. After addition of the elongation factors DSIF, PAF and SPT6 (75 pmol each), phosphorylation of EC* was performed with 0.4 μM P-TEFb and 1 mM ATP in 20 mM HEPES pH 7.4, 100 mM NaCl, 3 mM MgCl_2_ and 1 mM DTT for 30 min at 30 °C. The assembled EC* was kept on ice for 5 min and mixed with 3x molar excess of U1 snRNP (75 pmol) in a total volume of 37 μl. Following incubation on ice for 15 min, the sample was applied onto a Superose 6 Increase 3.2/300 column (GE Healthcare) equilibrated with the SEC100 buffer (20 mM HEPES pH 7.4, 100 mM NaCl, 3 mM MgCl_2_, 0.5 mM TCEP). Control runs of U1 snRNP alone and U1 snRNP with phosphorylated elongation factors were performed.

The fractions 1-12 were loaded individually onto 4-12% NuPAGE Bis-Tris gels (the amount loaded was calculated to have the same amount of Pol II in fraction #2) and run in 1x MOPS at 180 V for 50 min. Gels were stained with Instant Blue. To visualize U1 snRNA, fraction #2 of all samples as well as a positive control of U1 snRNP were incubated with 1 μl of proteinase K (NEB, 800 units/ml) for 30 min at 37 °C and loaded onto a denaturing gel (8% acrylamide/bis-acrylamide, 7M urea and 1x TBE). The denaturing gel was stained with SybrGold (Thermo Fisher Scientific) and visualized using a Typhoon 9500 FLA Imager (GE Healthcare). The U1-70k bands were confirmed by Western blotting using an antibody against U1-70k (Santa Cruz Biotechnology, sc-390899) and a secondary anti-mouse HRP antibody (Abcam, ab5870). The loading control was confirmed with an antibody against RPB3 (BETHYL, A303-771A) and a secondary anti-rabbit HRP antibody (GE Healthcare, NA934).

### RNA extension assay

The RNA extension assay was performed with a complementary DNA scaffold that corresponds to modified *JUNB* sequence with the 5’SS sequence inserted (in bold). Template DNA: 5’-TCT GGC GCG ATA GCT TTC CTG GCG TCG AAA CAC CCA GCA CCC AGC ACC CAG CAG GCA CCG **ATA CTT ACC TGG** TCC GCT CTC AAT GG-3’. Non-template DNA: 5’-CCA TTG AGA GCG GAC **CAG GTA AGT ATC** GGT GCC TGC TGG GTG CTG GGT GCT GGG TGT TTC GAC GCC AGG AAA GCT ATC GCG CCA GA-3’. RNA: 5’-/6-FAM/-UUU UUU CCA GGU AAG-3’. The scaffold has 14 nucleotides of upstream DNA, 63 nucleotides of downstream DNA, a 9 base-paired RNA·DNA hybrid and 6 nucleotides of exiting RNA. The oligos were synthesized by IDT.

RNA and template DNA were mixed in equimolar ratio (10 μM) in 20 mM HEPES pH 7.4, 100 mM NaCl and 3 mM MgCl_2_, and annealed by heating up at 60 °C for 4 min followed by decreasing the temperature by 1 °C min^-1^ steps to a final temperature of 30 °C in a thermocycler. All concentrations refer to the final concentration in the assays. *S. scrofa* Pol II (150 nM) and the RNA–template hybrid (100 nM) were incubated for 10 min at 30 °C, followed by addition of the non-template DNA (200 nM) and incubation at 30 °C for another 10 min. The phosphorylation reaction was then performed with 100 nM P-TEFb and 1 mM ATP in 20 mM HEPES pH 7.4, 100 mM NaCl, 3 mM MgCl_2_, 4% glycerol and 1 mM DTT for 15 min at 30 °C for either Pol II alone or in the presence of elongation factors (DSIF, PAF, SPT6 and RTF1, 150 nM each). The reactions were then cooled down on ice and a titration of U1 snRNP from 0 to 1.5 μM was added. Transcription was initiated upon addition of 50 μM NTP (Pol II alone) or 10 μM NTP (EC*) at 20 °C. The reactions were stopped at various timepoints by mixing 5 μl of transcription reaction with 5 μl of 2x stop buffer (6.4 M urea, 50 mM EDTA pH 8.0, 1x TBE). Samples were treated with 2 μl of proteinase K (NEB, 800 units/ml) for 20 min at 37 °C and 5 ul of each sample was applied onto 20% denaturing gels (20% acrylamide/bis-acrylamide, 7 M urea and 1xTBE, run in 0.5x TBE at 300 V for 100 min). The gels were visualized by the 6-FAM label on the RNA primer using the Typhoon 9500 FLA Imager (GE Healthcare).

### Cryo-EM data collection and processing

Cryo-EM data were collected on the 300 kV FEI Titan Krios with a K3 summit direct detector (Gatan) and a GIF quantum energy filter (Gatan) operated with a slit width of 20 eV. Automated data collection was performed with SerialEM (*41*) with a tilt angle of 40° at a nominal magnification of 81,000x, corresponding to a pixel size of 1.05 Å/pixel. Image stacks of 40 movie frames were collected with a defocus range of −0.5 to −2.0 μm in electron counting mode and a dose rate of 1.01 e^-^/Å^2^/frame. A total of 17,321 image stacks were collected across four datasets for the Pol II-U1 snRNP complex with 145-nt RNA. A total of 5,660 image stacks were collected for the Pol II-U1 snRNP complex with 155-nt RNA.

All movie frames were aligned and the contrast transfer function (CTF) parameters were calculated in Warp (*42*). Particles in 400 pixels x 400 pixels were selected by automatic particle picking in Warp. The following steps were performed in RELION 3.0 (*43*) to exclude bad particles from the dataset: 1) Two-dimensional (2D) classification was performed and particles in bad classes with poorly recognizable features were excluded. 2) In the second round of 2D classification, free U1 snRNP particles were excluded, leaving only Pol II containing particles. 3) The remaining particles were refined using three-dimensional (3D) refinement, followed by multiple rounds of CTF refinement to correct the CTF parameters. Beam-induced particle motion was corrected using Bayesian polishing in RELION 3.0 (*44, 45*). The particles were then refined with a soft mask on Pol II and divided into six classes using 3D classification in RELION with local fine-angle search (0.9 degree) (fig. S2, A and B). All 3D classes with bad particles or the class with missing clamp were discarded.

For the dataset of the Pol II-U1 snRNP complex with 145-nt RNA, all good particles with Pol II were combined and 3D refined using a soft mask on Pol II, resulting in a reconstruction of Pol II at 2.6 Å resolution. To separate Pol II alone particles from the Pol II-U1 snRNP particles, signal subtraction of Pol II followed by focused 3D classification of the subtracted particles without alignment was performed using a large spherical mask near the RNA exit site (where the extra density is) (fig. S2, C and D). The classes with an extra density corresponding to U1 snRNP were combined (17.5%), reverted to original particles and 3D refined with a solvent mask. This resulted in a Pol II-U1 snRNP reconstruction with 220,922 particles at an overall resolution of 3.0 Å (fig. S2E). A second round of signal subtraction and focused 3D classification without alignment using a soft mask on U1-70k/stem loop I (the interface density) was performed to separate the movements of U1 snRNP (fig. S2G). All four classes have good densities corresponding to U1-70k/stem loop I, yet a clear movement relative to Pol II was observed when they were superimposed. The class with the best density for U1-70k/stem loop I was selected, reverted to original particles and 3D refined with a solvent mask. This resulted in the final Pol II-U1 snRNP reconstruction with 61,596 particles at an overall resolution of 3.6 Å (fig. S2H).

To improve local resolutions for model building, the particles were also refined with a soft mask on either Pol II or the Pol II-U1 snRNP interface region (3.3 Å and 3.2 Å, respectively, fig. S2I). Following 3D refinement with a soft mask on Pol II, signal subtraction of Pol II was performed, leaving only the signal of U1 snRNP. The subtracted particles were then refined with a soft mask on U1 snRNP, resulting in a U1 snRNP reconstruction of 5.9 Å (fig. S2I). Due to the extended structure of U1 snRNP, the Sm-ring and RNA stem-loops (SL) 2-4 that are further away from the Pol II-U1 snRNP interface showed large movements relative to Pol II and their fit relied on the focused refined map of U1 snRNP (fig. S3 and fig. S4).

The dataset for the Pol II-U1 snRNP complex with 155-nt RNA was processed in the same way, resulting in a final reconstruction with 14,497 particles at 3.9 Å resolution (fig. S8). The same protein interface between Pol II and U1 snRNP was observed, whereas the linking RNA showed higher mobility and more diffuse densities.

Final maps were sharpened using PHENIX AutoSharpen (*46*). Local resolution of the maps was estimated using RELION 3.0 (*43*). All resolution estimations were based on the gold-standard Fourier Shell Correlation (FSC) calculations using the FSC = 0.143 criterion. A summary of all EM reconstructions obtained in this paper is listed in table S1.

### Model building and refinement

Initial models of *S. scrofa* Pol II (PDB: 6GMH (*27*)) and human U1 snRNP (PDB: 3PGW (*28*), 4PJO and 4PKD (*29*)) were rigid-body fitted into the overall map in Chimera (*47*). The Pol II model was manually adjusted in Coot (*48*) using the high-resolution map of Pol II (2.6 Å) and the structure was real-space refined in PHENIX (*49*). The refined Pol II model was then fitted into the overall Pol-U1 snRNP map and adjusted in Coot. The U1 snRNP model was first rebuilt in the focused refined map of U1 snRNP in Coot and the structure was real-space refined in PHENIX and used later as a restraint. Both Pol II and U1 snRNP models were fitted into the interface focused refined map, and the models for RPB2, RPB12, U1-70k and stem loop I of U1 snRNA were rebuilt in Coot. The resulting complete model of Pol II-U1 snRNP was then real-space refined in the overall map using PHENIX and structure restraints of U1 snRNP.

We could not observe density for the exposed Pol II stalk RPB4/RPB7, as it is often the case for Pol II structures. In addition, no density corresponding to U1A was observed, likely due to the high flexibility of the stem loop II of U1 snRNP. We therefore modelled the Pol II stalk and U1A by superposition of previous structures of Pol II and U1 snRNP (PDB: 6GMH (*27*), 3PGW (*28*), 4PJO and 4PKD (*29*)). We also did not observe any density for U1C.

Figures were generated using PyMOL (The PyMOL Molecular Graphics System, Version 2.0 Schrödinger, LLC.), Chimera (*47*) and Chimera X (*50*). Sequence alignment was performed using Jalview (*51*).

---

**Fig. S1.**
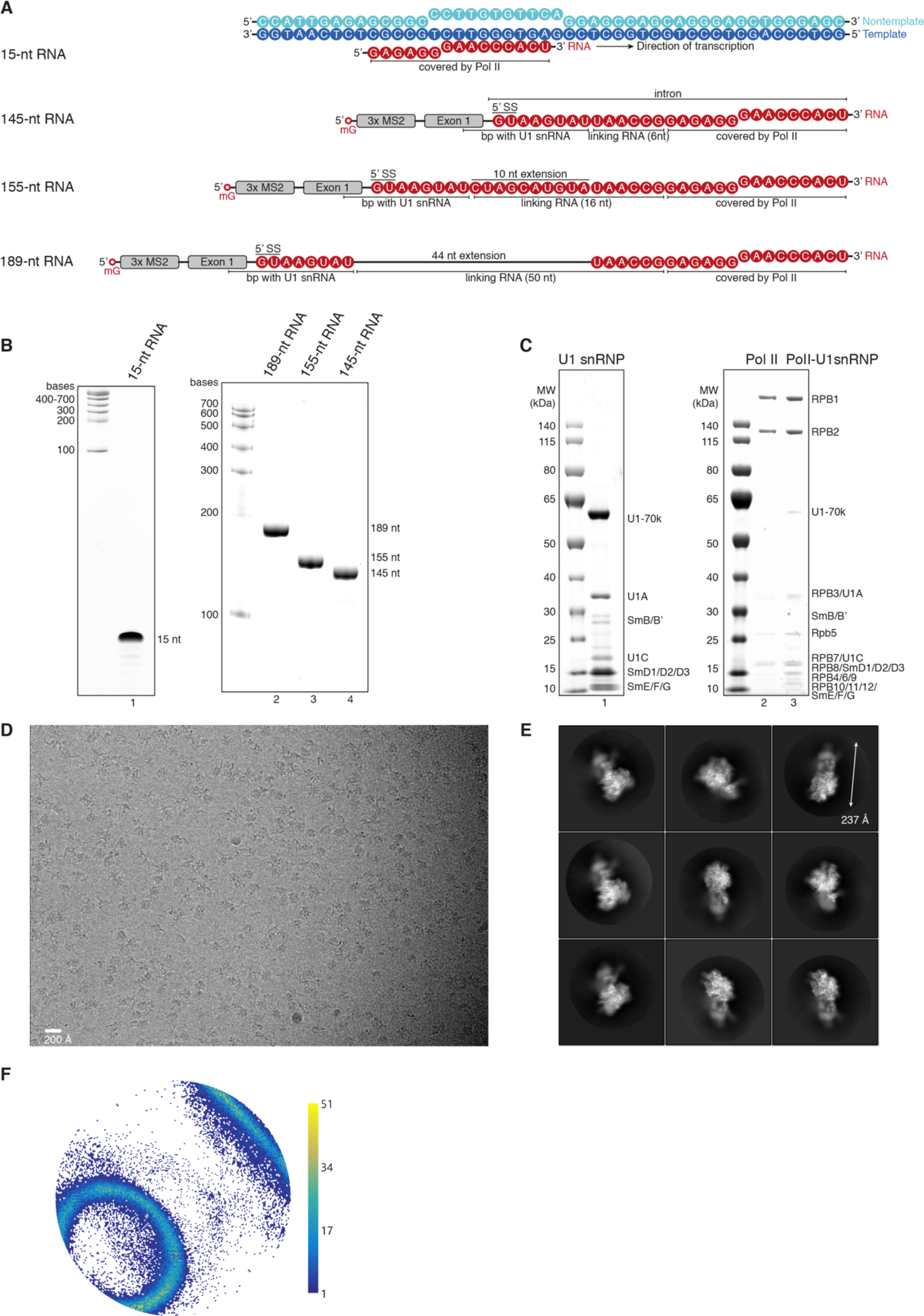
Preparation of the transcribing Pol II-U1 snRNP complex. (**A**) Schematic of the DNA scaffold and the RNA constructs used for structural and biochemical analysis. The 15-nt RNA is designed to be completely covered within Pol II. The 145-nt RNA consists of three MS_2_-tags, a 5’ exon and a truncated intron of 29 nt, of which 6 nt (linking RNA) are exposed between Pol II and U1 snRNP. The 155-nt RNA contains an extension of 10 nt within the linking RNA, while the 189-nt RNA contains a linking RNA of 50 nt, sufficient for assembly of the B complex. (**B**) Purified RNAs on a 20% denaturing gel (15-nt RNA, lane 1) or an 8% denaturing gel (145-nt, 155-nt and 189-nt RNAs), stained with Toluidine Blue. (**C**) Purified human U1 snRNP and *S. scrofa* Pol II on 4-12% NuPAGE Bis-Tris gels run in MOPS, stained with Instant Blue. The Pol II-U1 snRNP on lane 3 was used for freezing grids. (**D**) Representative micrograph collected on the FEI 300 kV Titan Krios with a K3 detector in electron counting mode and a 40° tilt. (**E**) Two-dimensional averages of the transcribing Pol II-U1 snRNP complex containing the 145-nt RNA, with a dimension of 237 Å across Pol II and U1 snRNP. (**F**) Angular distribution plot of the overall map, with the scale showing the number of particles assigned to a particular angle. The ring-shape of the angular distribution is resulted from the tilted data collection to resolve the preferred orientation of the complex.

**Fig. S2.**
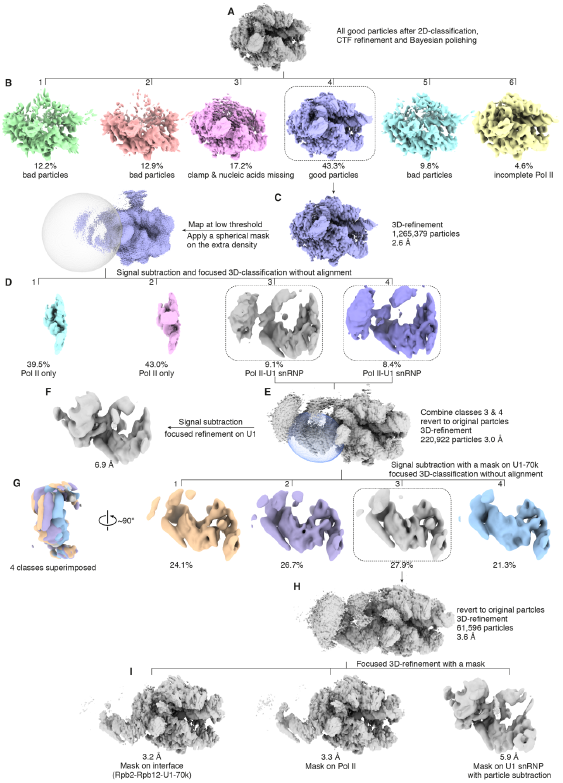
Cryo-EM data processing. After removing bad particles and U1 snRNP alone particles with two-dimensional classification, particles were combined and 3D-refined, followed by CTF refinement and Baysian polishing in RELION 3.0 (**A**). The particles were subsequently divided into six classes by 3D-classification with a fine-angle local search and the remaining bad particles as well as incomplete Pol II particles were removed (**B**). Good particles were 3D-refined to obtain a high-resolution map of Pol II (**C**). Following signal subtraction of Pol II and focused 3D-classification without alignment of the subtracted particles with a large spherical soft mask onto the extra density, the classes containing densities corresponding to U1 snRNP were selected and combined (**D**). This resulted in a reconstruction of Pol II-U1 snRNP at 3.0 Å resolution (**E**), with a resolution of 6.9 Å for U1 snRNP (**F**). The movement of U1 snRNP relative to Pol II was further separated by signal subtraction of Pol II and focused 3D-classification without alignment with a soft spherical mask on U1-70k (**G**). The final reconstruction has an overall resolution of 3.6 Å (**H**) and focused 3D-refinement on Pol II, the Pol II-U1 snRNP interface and U1 snRNP improved the resolution of respective regions and aided model building (**I**).

**Fig. S3.**
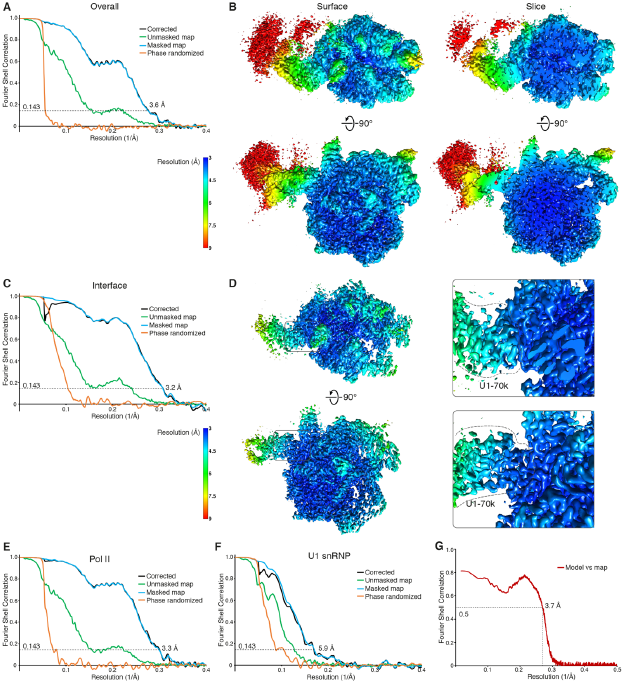
Local resolution maps and FSC curves. (**A**) Gold-standard Fourier Shell Correlation (FSC) curves of the overall map of Pol II-U1 snRNP. (**B**) Local resolution estimations of the overall map in both surface and slice views showed that the Pol II core is beyond 3 Å (deep blue), while the Pol II-U1 snRNP interface is around 4-4.5 Å (cyan) and the rest of U1 snRNP in the range of 6-9 Å (green-red). (**C**) FSC curves of the focused refined map at the Pol II-U1 snRNP interface. (**D**) Local resolution estimation of the interface focused refined map showed improved densities for U1-70k RRM at the interface (3-4 Å), where densities for bulky side chains could be observed. (E and F) FSC curves of the focused refined maps on Pol II (**E**) and U1 snRNP (**F**), respectively. (**G**) Model versus map FSC for the overall map using the FSC standard of 0.5.

**Fig. S4.**
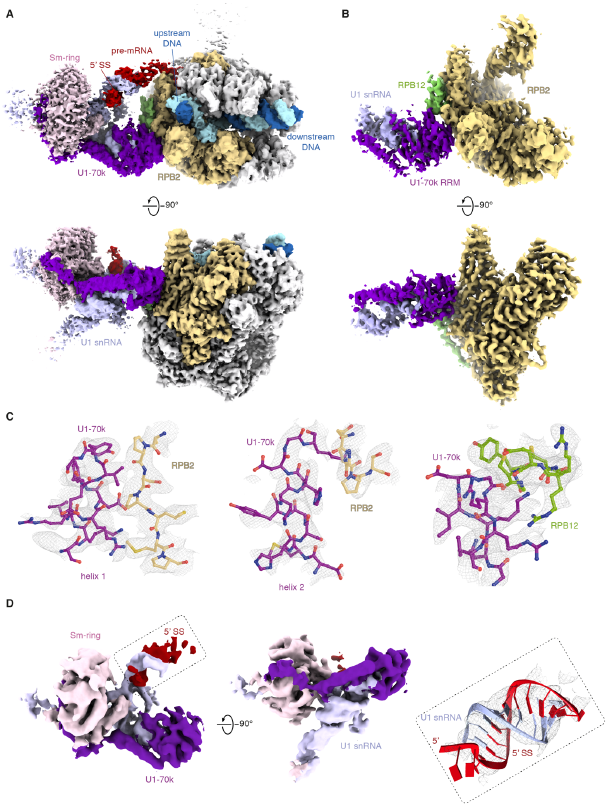
Cryo-EM densities of Pol II-U1 snRNP. (**A**) Cryo-EM densities of the overall map, where weak densities were observed for the linking RNA and the Pol II stalk. The views and colors are the same as in Fig. 1, B and C. (**B**) Cryo-EM densities of the focused refined map at the interface region, with densities for U1-70k colored in purple, RPB2 in gold and RPB12 in green. (**C**) Cryo-EM densities at the Pol II-U1 snRNP interface shown in grey mesh with models of U1-70k, RPB2 and RPB12 in cartoon. Densities for bulky side chains at the interface were clearly resolved. (**D**) Cryo-EM densities of the focused refined map for U1 snRNP in two views, with improved densities for U1 snRNA and Sm-ring. Densities for the 5’SS-U1 snRNA duplex are shown in grey mesh with the RNA model fitted.

**Fig. S5.**
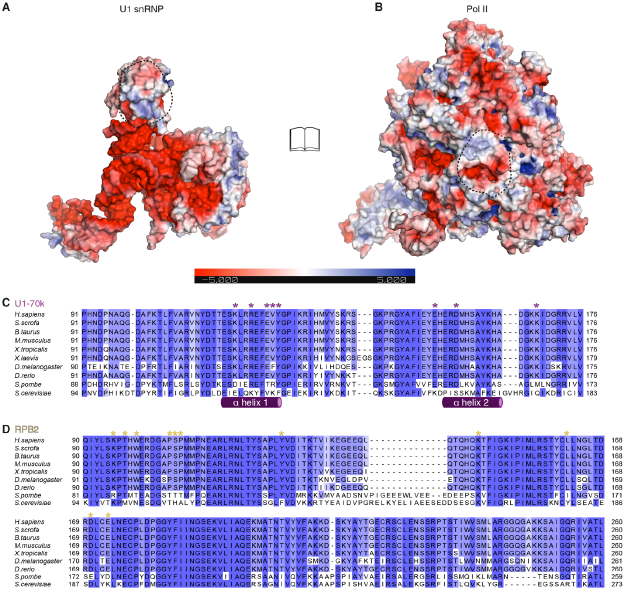
Electrostatic surface potentials and sequence alignments. (**A** and **B**) The electrostatic surface potential of U1 snRNP (A) and Pol II (B) in book view with a range of ± 5 kT/e, where deep blue represents positively and red represents negatively charged areas. The dashed lines highlight the interface areas between Pol II and U1 snRNP that are oppositely charged. (**C**) Sequence alignment of U1-70k RRM from yeast to human. The key Pol II interacting residues are marked with purple stars. (**D**) Sequence alignment of the RPB2 region that contacts U1-70k RRM from yeast to human. The key U1-70k interacting residues are marked with golden stars.

**Fig. S6.**
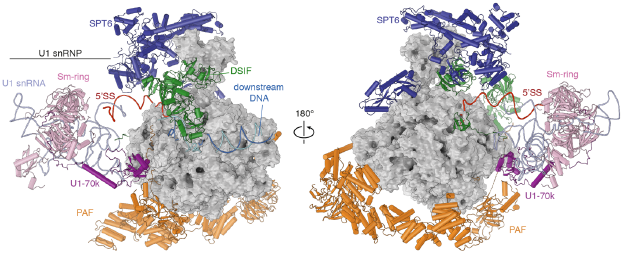
Superposition with activated elongation complex EC*. Superposition of the Pol II-U1 snRNP complex with the activated transcription elongation complex EC* (PDB: 6GMH (*27*)). Pol II is shown in surface representation in grey, while U1 snRNP is colored in pink except for U1-70k in purple. Template DNA is depicted as blue ribbon and non-template DNA in cyan. RNA is shown as red ribbon. The elongation factors SPT6 (dark blue), DSIF (green) and PAF (orange) are compatible with U1 snRNP on the Pol II surface. The view in the left panel is the same as Fig. 1B.

**Fig. S7.**
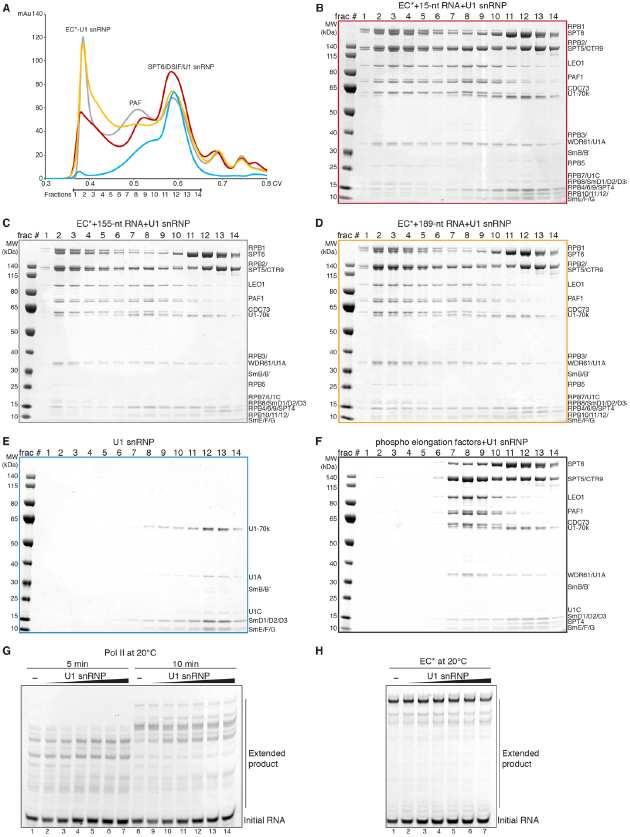
Analytical size exclusion assay for EC*-U1 snRNP complex formation and RNA extension assays. (**A**) Size exclusion chromatograms of complex formation between the activated transcription elongation complex (EC*) and U1 snRNP using three different RNA constructs. Profiles for EC*-U1 snRNP complex formation with 15-nt RNA (red), 155-nt RNA (grey) and 189-nt RNA (orange) as well as a control profile of U1 snRNP alone (blue) are shown. Fractions used for analysis on SDS-PAGE are marked. (**B to F**) Fractions of the analytical size exclusion assays were analyzed on a 4-12% NuPAGE Bis-Tris gel for EC*-U1 snRNP complex formation with 15-nt RNA (B), 155-nt RNA (C) and 189-nt RNA (D), as well as for the control runs with U1 snRNP alone (E) and U1 snRNP together with phosphorylated elongation factors (F). Both the 155-nt and 189-nt RNAs enabled EC*-U1 snRNP complex formation. The minimal 15-nt RNA that lacks the 5’SS still allowed U1 snRNP binding, although at apparently sub-stoichiometric levels. (**G** and **H**) RNA extension assays showed that U1 snRNP binding does not affect the transcription speed for either phosphorylated Pol II alone (G) or EC* (H). A titration of U1 snRNP from 0 to 10x molar excess to Pol II was used. All assays were repeated in triplicate.

**Fig. S8.**
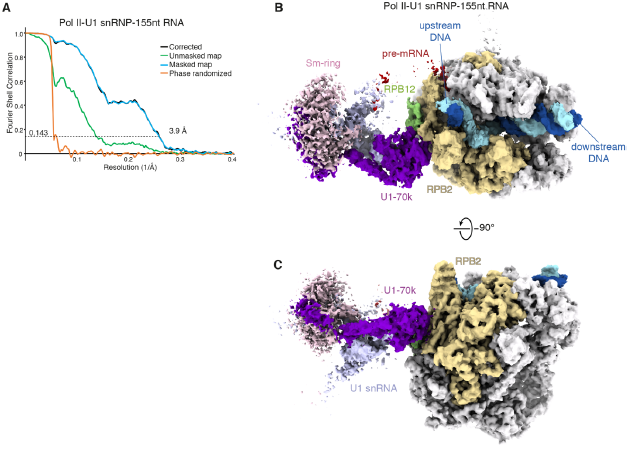
Cryo-EM analysis of alternative Pol II-U1 snRNP complex with 155-nt RNA. (**A**) FSC curves of the overall map of the Pol II-U1 snRNP complex with 155-nt RNA (10-nt extension in the linking RNA). (**B** and **C**) Cryo-EM densities of the Pol II-U1 snRNP complex with 155-nt RNA, where the linking RNA became more mobile compared to 145-nt RNA. The same protein interface among U1-70k, RPB2 and RPB12 was observed. The views and colors are the same as in Fig. 1, B and C, and fig. S4A.

**Fig. S9.**
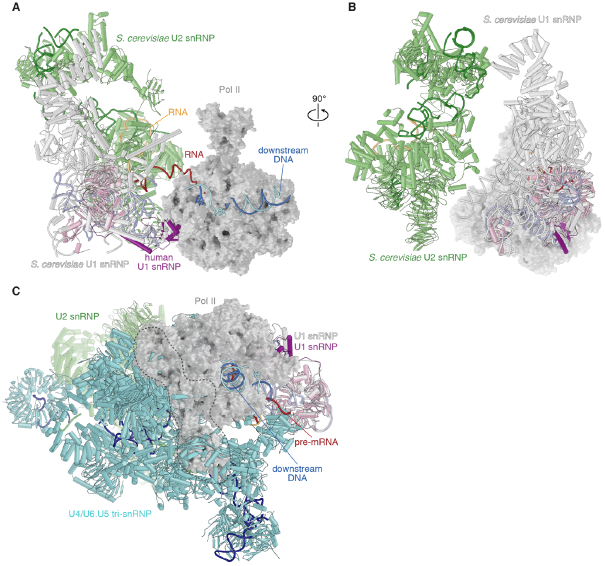
Superposition with spliceosome structures. (**A** and **B**) Superposition of the mammalian Pol II-U1 snRNP structure with the yeast pre-spliceosome or A complex (PDB: 6G90 (*34*)) on the U1 snRNP Sm-ring revealed that A complex formation can happen on the surface of Pol II (grey surface). *S. cerevisiae* U1 snRNP is in white and *S. cerevisiae* U2 snRNP in light green. The pre-mRNA of the Pol II-U1 snRNP complex is colored in red, while that of the yeast A complex in orange. (**C**) Superposition of the Pol II-U1 snRNP structure with the human pre-B complex (PDB: 6QX9 (*31*)) on the U1 snRNP Sm-ring showed some clashes (highlighted with dashed lines) between Pol II (grey surface) and tri-snRNP (cyan). However, the flexibility of the pre-B complex may enable structural rearrangements (compare main text).

**Table S1.** Statistics of cryo-EM reconstructions and structural model.

|  | Transcribing Pol II-U1 snRNP complex |  |  |  |
| --- | --- | --- | --- | --- |
|  | 145-nt RNA |  |  | 155-nt RNA |
|  | Overall<br>(EMD-xxxx) | Interface<br>(EMD-xxxx) | U1 snRNP<br>(EMD-xxxx) | Overall<br>(EMD-xxxx) |
| Data collection and processing |  |  |  |  |
| Magnification | 81 000 x |  |  | 81 000 x |
| Voltage (kV) | 300 |  |  | 300 |
| Electron exposure (e<sup>-</sup>/Å<sup>2</sup>) | 40.4 |  |  | 40.4 |
| Defocus range (μm) | 0.5-2.0 |  |  | 0.5-2.0 |
| Pixel size (Å) | 1.05 |  |  | 1.05 |
| Symmetry imposed | C1 |  |  | C1 |
| Initial particle numbers | 1,265,379 |  |  | 320,847 |
| Final particle numbers | 61,596 |  |  | 14,497 |
| Map resolution (Å) | 3.6 | 3.2 | 5.9 | 3.9 |
| Overall B-sharpen applied | -72.73 | -69.66 | -360.19 | -76.06 |
| Refinement |  |  |  |  |
| Initial models used (PDB code) | 6GMH, 3PGW,<br>4PJO, 4PKD |  |  |  |
| Model resolution (Å) | 3.7 |  |  |  |
| Model composition |  |  |  |  |
| Non-hydrogen atoms | 43,792 |  |  |  |
| Protein residues | 4,745 |  |  |  |
| Nucleic acid residues | 274 |  |  |  |
| Ligands | Zn:8 Mg:1 |  |  |  |
| Mean B factors (Å<sup>2</sup>) |  |  |  |  |
| Protein | 74.31 |  |  |  |
| Nucleotide | 230.98 |  |  |  |
| Ligand | 72.36 |  |  |  |
| R.m.s. deviations |  |  |  |  |
| Bond lengths (Å) | 0.002 |  |  |  |
| Bond angles (°) | 0.416 |  |  |  |
| Validation |  |  |  |  |
| MolProbity score | 1.23 |  |  |  |
| Clash score | 4.27 |  |  |  |
| Poor rotamers (%) | 0.02 |  |  |  |
| Ramachandran plot |  |  |  |  |
| Favored (%) | 97.89 |  |  |  |
| Allowed (%) | 2.11 |  |  |  |
| Outliers (%) | 0.0 |  |  |  |
| PDB code | XXXX |  |  |  |

## References

1. K. M. Kotovic, D. Lockshon, L. Boric, K. M. Neugebauer, Cotranscriptional recruitment of the U1 snRNP to intron-containing genes in yeast. Mol Cell Biol 23, 5768–5779 (2003).

2. S. A. Lacadie, M. Rosbash, Cotranscriptional spliceosome assembly dynamics and the role of U1 snRNA:5’ss base pairing in yeast. Mol Cell 19, 65–75 (2005).

3. I. Listerman, A. K. Sapra, K. M. Neugebauer, Cotranscriptional coupling of splicing factor recruitment and precursor messenger RNA splicing in mammalian cells. Nat Struct Mol Biol 13, 815–822 (2006).

4. E. W. J. Wallace, J. D. Beggs, Extremely fast and incredibly close: cotranscriptional splicing in budding yeast. RNA 23, 601–610 (2017).

5. G. Bird, D. A. Zorio, D. L. Bentley, RNA polymerase II carboxy-terminal domain phosphorylation is required for cotranscriptional pre-mRNA splicing and 3’-end formation. Mol Cell Biol 24, 8963–8969 (2004).

6. D. L. Bentley, Coupling mRNA processing with transcription in time and space. Nat Rev Genet 15, 163–175 (2014).

7. M. Tellier, I. Maudlin, S. Murphy, Transcription and splicing: A two-way street. Wiley Interdiscip Rev RNA 11, e1593 (2020).

8. L. Herzel, D. S. M. Ottoz, T. Alpert, K. M. Neugebauer, Splicing and transcription touch base: co-transcriptional spliceosome assembly and function. Nat Rev Mol Cell Biol 18, 637–650 (2017).

9. L. Wachutka, L. Caizzi, J. Gagneur, P. Cramer, Global donor and acceptor splicing site kinetics in human cells. Elife 8, (2019).

10. P. J. Curtis, N. Mantei, C. Weissmann, Characterization and kinetics of synthesis of 15S beta-globin RNA, a putative precursor of beta-globin mRNA. Cold Spring Harb Symp Quant Biol 42 Pt 2, 971–984 (1978).

11. A. Audibert, D. Weil, F. Dautry, In vivo kinetics of mRNA splicing and transport in mammalian cells. Mol Cell Biol 22, 6706–6718 (2002).

12. M. K. Sakharkar, V. T. Chow, P. Kangueane, Distributions of exons and introns in the human genome. In Silico Biol 4, 387–393 (2004).

13. M. J. Dye, N. Gromak, N. J. Proudfoot, Exon tethering in transcription by RNA polymerase II. Mol Cell 21, 849–859 (2006).

14. L. E. Giono, A. R. Kornblihtt, Linking transcription, RNA polymerase II elongation and alternative splicing. Biochem J 477, 3091–3104 (2020).

15. M. de la Mata et al., A slow RNA polymerase II affects alternative splicing in vivo. Mol Cell 12, 525–532 (2003).

16. A. Pandya-Jones, D. L. Black, Co-transcriptional splicing of constitutive and alternative exons. RNA 15, 1896–1908 (2009).

17. G. Dujardin et al., How slow RNA polymerase II elongation favors alternative exon skipping. Mol Cell 54, 683–690 (2014).

18. A. Yuryev et al., The C-terminal domain of the largest subunit of RNA polymerase II interacts with a novel set of serine/arginine-rich proteins. Proc Natl Acad Sci U S A 93, 6975–6980 (1996).

19. T. Nojima et al., Mammalian NET-Seq Reveals Genome-wide Nascent Transcription Coupled to RNA Processing. Cell 161, 526–540 (2015).

20. T. Misteli, D. L. Spector, RNA polymerase II targets pre-mRNA splicing factors to transcription sites in vivo. Mol Cell 3, 697–705 (1999).

21. K. M. Harlen et al., Comprehensive RNA Polymerase II Interactomes Reveal Distinct and Varied Roles for Each Phospho-CTD Residue. Cell Rep 15, 2147–2158 (2016).

22. C. J. David, A. R. Boyne, S. R. Millhouse, J. L. Manley, The RNA polymerase II C-terminal domain promotes splicing activation through recruitment of a U2AF65-Prp19 complex. Genes Dev 25, 972–983 (2011).

23. B. J. Natalizio, N. D. Robson-Dixon, M. A. Garcia-Blanco, The Carboxyl-terminal Domain of RNA Polymerase II Is Not Sufficient to Enhance the Efficiency of Pre-mRNA Capping or Splicing in the Context of a Different Polymerase. J Biol Chem 284, 8692–8702 (2009).

24. C. L. Will, R. Luhrmann, Spliceosome structure and function. Cold Spring Harb Perspect Biol 3, (2011).

25. S. M. Mount, I. Pettersson, M. Hinterberger, A. Karmas, J. A. Steitz, The U1 small nuclear RNA-protein complex selectively binds a 5’ splice site in vitro. Cell 33, 509–518 (1983).

26. C. Boesler et al., A spliceosome intermediate with loosely associated tri-snRNP accumulates in the absence of Prp28 ATPase activity. Nat Commun 7, 11997 (2016).

27. S. M. Vos et al., Structure of activated transcription complex Pol II-DSIF-PAF-SPT6. Nature 560, 607–612 (2018).

28. G. Weber, S. Trowitzsch, B. Kastner, R. Luhrmann, M. C. Wahl, Functional organization of the Sm core in the crystal structure of human U1 snRNP. EMBO J 29, 4172–4184 (2010).

29. Y. Kondo, C. Oubridge, A. M. van Roon, K. Nagai, Crystal structure of human U1 snRNP, a small nuclear ribonucleoprotein particle, reveals the mechanism of 5’ splice site recognition. Elife 4, (2015).

30. X. Zhan, C. Yan, X. Zhang, J. Lei, Y. Shi, Structures of the human pre-catalytic spliceosome and its precursor spliceosome. Cell Res 28, 1129–1140 (2018).

31. C. Charenton, M. E. Wilkinson, K. Nagai, Mechanism of 5’ splice site transfer for human spliceosome activation. Science 364, 362–367 (2019).

32. B. Spiluttini et al., Splicing-independent recruitment of U1 snRNP to a transcription unit in living cells. J Cell Sci 123, 2085–2093 (2010).

33. K. Bertram et al., Cryo-EM Structure of a Pre-catalytic Human Spliceosome Primed for Activation. Cell 170, 701-713 e711 (2017).

34. C. Plaschka, P. C. Lin, C. Charenton, K. Nagai, Prespliceosome structure provides insights into spliceosome assembly and regulation. Nature 559, 419–422 (2018).

35. S. Sharma, S. P. Wongpalee, A. Vashisht, J. A. Wohlschlegel, D. L. Black, Stem-loop 4 of U1 snRNA is essential for splicing and interacts with the U2 snRNP-specific SF3A1 protein during spliceosome assembly. Genes Dev 28, 2518–2531 (2014).

36. S. Vagner, U. Ruegsegger, S. I. Gunderson, W. Keller, I. W. Mattaj, Position-dependent inhibition of the cleavage step of pre-mRNA 3’-end processing by U1 snRNP. RNA 6, 178–188 (2000).

37. D. Kaida et al., U1 snRNP protects pre-mRNAs from premature cleavage and polyadenylation. Nature 468, 664–668 (2010).

38. A. E. Almada, X. Wu, A. J. Kriz, C. B. Burge, P. A. Sharp, Promoter directionality is controlled by U1 snRNP and polyadenylation signals. Nature 499, 360–363 (2013).

39. P. K. Andersen, S. Lykke-Andersen, T. H. Jensen, Promoter-proximal polyadenylation sites reduce transcription activity. Genes Dev 26, 2169–2179 (2012).

40. M. Zillmann, M. L. Zapp, S. M. Berget, Gel electrophoretic isolation of splicing complexes containing U1 small nuclear ribonucleoprotein particles. Mol Cell Biol 8, 814–821 (1988).

41. D. N. Mastronarde, Automated electron microscope tomography using robust prediction of specimen movements. J Struct Biol 152, 36–51 (2005).

42. D. Tegunov, P. Cramer, Real-time cryo-electron microscopy data preprocessing with Warp. Nat Methods 16, 1146–1152 (2019).

43. S. H. Scheres, Processing of Structurally Heterogeneous Cryo-EM Data in RELION. Methods Enzymol 579, 125–157 (2016).

44. S. H. Scheres, A Bayesian view on cryo-EM structure determination. J Mol Biol 415, 406–418 (2012).

45. S. H. Scheres, RELION: implementation of a Bayesian approach to cryo-EM structure determination. J Struct Biol 180, 519–530 (2012).

46. T. C. Terwilliger, O. V. Sobolev, P. V. Afonine, P. D. Adams, Automated map sharpening by maximization of detail and connectivity. Acta Crystallogr D Struct Biol 74, 545–559 (2018).

47. E. F. Pettersen et al., UCSF Chimera--a visualization system for exploratory research and analysis. J Comput Chem 25, 1605–1612 (2004).

48. P. Emsley, B. Lohkamp, W. G. Scott, K. Cowtan, Features and development of Coot. Acta Crystallogr D Biol Crystallogr 66, 486–501 (2010).

49. D. Liebschner et al., Macromolecular structure determination using X-rays, neutrons and electrons: recent developments in Phenix. Acta Crystallogr D Struct Biol 75, 861–877 (2019).

50. T. D. Goddard et al., UCSF ChimeraX: Meeting modern challenges in visualization and analysis. Protein Sci 27, 14–25 (2018).

51. A. M. Waterhouse, J. B. Procter, D. M. Martin, M. Clamp, G. J. Barton, Jalview Version 2--a multiple sequence alignment editor and analysis workbench. Bioinformatics 25, 1189–1191 (2009).

